# Identification and functional investigation of *Octopus vulgaris* TRPV channels as potential nociceptors in cephalopods

**DOI:** 10.64898/2026.03.27.714695

**Authors:** Eleonora Maria Pieroni, Howard Baylis, Vincent O’Connor, Lindy Holden-Dye, Luis Alfonso Yañez-Guerra, Pamela Imperadore, Graziano Fiorito, James Dillon

## Abstract

Nociception is an essential response for organisms to avoid potential harm and promote survival. Its molecular determinants are largely conserved across Eumetazoa. TRPV receptors are polymodal ion channels exhibiting selective peripheral expression and functional coupling that underpins nociception and pain modulation in complex organisms. However, the execution of protective behaviours triggered by TRPVs is also found in species with a simpler nervous organisation, thus encouraging their investigation in invertebrate model organisms to increase understanding of animal nociception.

Cephalopods represent an interesting invertebrate phylum with respect to the evolution of the nervous system, whose complexity suggests it might support pain-like states that exist in vertebrates. This possibility is reflected by the inclusion of cephalopods in the UK and EU animal welfare legislations. Despite this, there is poor characterisation of cephalopod molecular nociceptors.

For this reason, we used *in silico* analysis to identify two TRPV channels in *Octopus vulgaris* genome (*Ovtrpv1* and *Ovtrpv2*). We validated the putative transcript sequences and highlighted prevalent expression in sensory tissues. We investigated the functional competence of these TRPVs by heterologously expressing *Ovtrpv1* and *Ovtrpv2* cDNA into *Caenorhabditis elegans* null mutants of the orthologous genes, *ocr-2* and *osm-9* respectively. *Ovtrpvs* successfully rescued the aversive response to chemical and mechanical noxious stimuli in the *C. elegans* mutants, suggesting these receptors are polymodal nociceptors. Additionally, complementary investigation using *Xenopus laevis* oocytes showed *Ovtrpv1* and *Ovtrpv2* form an active heteromeric channel gated by nicotinamide. This study highlights *Ovtrpvs* as an important route to better understand nociceptive detection in cephalopods.

## Background

General nociception executes function by activating a reflexive withdrawal response, protecting organisms from potentially threatening stimuli. Various avoidance behaviours operate across animal phyla and advantages organismal survival (Smith and Lewin, 2009). Nociception is characterised by a conserved specialised neuronal architecture and shared molecular determinants that persist across distinct nervous system organisations and largely encompasses a core reflex that triggers withdrawal avoidance responses (Julius and Basbaum, 2001). In more complex organisms, however, the reflexive nociceptive responses are modulated by top-down neuronal signalling leading to the elaboration of learnt and emotionally encoded pain states (Basbaum *et al*., 2009).

Cephalopods are invertebrate species characterised by well described withdrawal responses following exposure to noxious challenges (Crook, Hanlon and Walters, 2013; Hague, Florini and Andrews, 2013; Crook, 2021). Furthermore, these molluscs possess a central nervous system that can act in a hierarchical fashion, allowing them to express complex behaviours (Amodio *et al*., 2019; Schnell *et al*., 2021). The details of this anatomy are different from organisms that are unequivocally known to express the emotional states defined by pain, nonetheless cephalopods’ central brain mass complexity provokes the possibility that their neural architecture is organised to allow brain states with components that resemble pain (Andrews *et al*., 2013; Smith *et al*., 2013; Shigeno *et al*., 2018). This highlights potential ethical issues and associated welfare constraints about the experimental use of cephalopods that have seen these animals included in research legislations (UK S.I. 1993/2103, 1993; European Parliament and Union, 2010).

This raises the scientific value of detailing the biological organisation of nociception reflexes and their modulation in this important biological and societally utilised taxon.

In our previous study, we identified *O. vulgaris* orthologues of genes with a defined role in nociception and pain to enrich understanding of the molecular determinants of these phenomena in cephalopods. We coupled this to functional studies in *C. elegans* loss of function mutants of the corresponding genes of interest and identified 19 potential candidates warranting more detailed investigation for their role in *O. vulgaris* nociception (Pieroni *et al*., 2026). Among our candidates, the previously investigated vanilloid transient receptor potential (TRPV) emerged as a priority candidate for further investigation (Caterina *et al*., 1997).

TRP receptors are a large family of ion channels highly conserved across animal species due to their extensive role in different physiological functions (Pedersen, Owsianik and Nilius, 2005; Ramsey, Delling and Clapham, 2006; Venkatachalam and Montell, 2007). Among the seven major TRP subfamilies, the vanilloid sensitive homologues, include receptors that detect aversive stimuli (Colton, 2006; Radresa *et al*., 2012; Shibasaki, 2024). The best characterised representative of TRPVs in mammals is the capsaicin receptor TRPV1 (Caterina *et al*., 1997). TRPV1 is a polymodal nociceptor which, in addition to vanilloid compounds, responds to cell damaging pH and temperature changes (Venkatachalam and Montell, 2007; Dhaka *et al*., 2009; Julius, 2013). TRPVs are typically tetrameric membrane receptors in which individual subunits have six transmembrane domains (TMs), cytosolic N-and C-terminals and a hydrophobic short P-loop between the fifth and sixth TMs that creates the selectivity filter (Rosasco and Gordon, 2017). This structure and the sensory related functions it underpins, are conserved across vertebrate and invertebrate species. This includes *Hirudo medicinalis, C. elegans and Drosophila melanogaster*, where their contribution to nociception is well-documented (Colbert, Smith and Bargmann, 1997; Tobin *et al*., 2002; Gong *et al*., 2004; Summers, Holec and Burrell, 2014; Ohnishi *et al*., 2020).

In this study, we characterised two TRPV channel representatives (*Ovtrpv1* and *Ovtrpv2*) from *O. vulgaris.* These are orthologues of *C. elegans ocr* and *osm-9* receptors respectively, enabling us to confirm function by heterologous expression in *C. elegans* null mutants and successful rescue of the aversive response in these strains. We used *in silico* and experimental analyses encompassing PCR to localise *Ovtrpvs* expression to distinct tissues, and *Xenopus* oocytes expression to characterise channel function. Taken together these data validated two *bona fide* TRPV channel representatives from *O. vulgaris* which have the functional properties and anatomical localisation to act as polymodal sensory receptors with a key role in detecting and classifying environmental stimuli.

## Results

### *In silico* identification of *O. vulgaris trpv* transcript and its experimental validation

When interrogating the previously available *O. vulgaris* transcriptome (Petrosino, 2015; Petrosino *et al*., 2022) using sequences of curated TRPV channels (e.g., *H. sapiens* TRPV1), we retrieved two hits (c32354_g7_i1 and c32354_g6_i1). However, *in silico* translation of these sequences revealed that the retrieved hits corresponded to incomplete fragments of a larger transcript. We therefore performed an *in silico* analysis reiterating the sequences between *O. vulgaris, O. bimaculoides and Aplysia californica* transcriptome databases. This led to the identification of three additional overlapping fragments (c32354_g14_i1, c32354_g12_i1, c32354_g2_i1) generating a contig transcript encoding a putative full length TRPV channel subunit (transcript accession number: PV164572, Pieroni *et al.,* 2026). Secondary structure prediction tools identified intracellular N- and C-terminals, six TMs and several ankyrin repeats in the N-terminal region (Figure 1). Using 3D modelling with AlphaFold 3 (Abramson *et al*., 2024) and simulation of the protein structure within the membrane with PPM3 server (Lomize, Todd and Pogozheva, 2022), we revealed the presence of a P-loop between TM 5 and TM 6, another key signature of these channels that was not detectable with secondary structure prediction tools (Figure 1). We designed primers that flank the predicted full length sequence and used PCR amplification from *O. vulgaris* reverse transcribed mRNA combined with cDNA sequencing to authenticate it as a *bona fide*sequence (accession number: PX926345).

**Figure 1.**
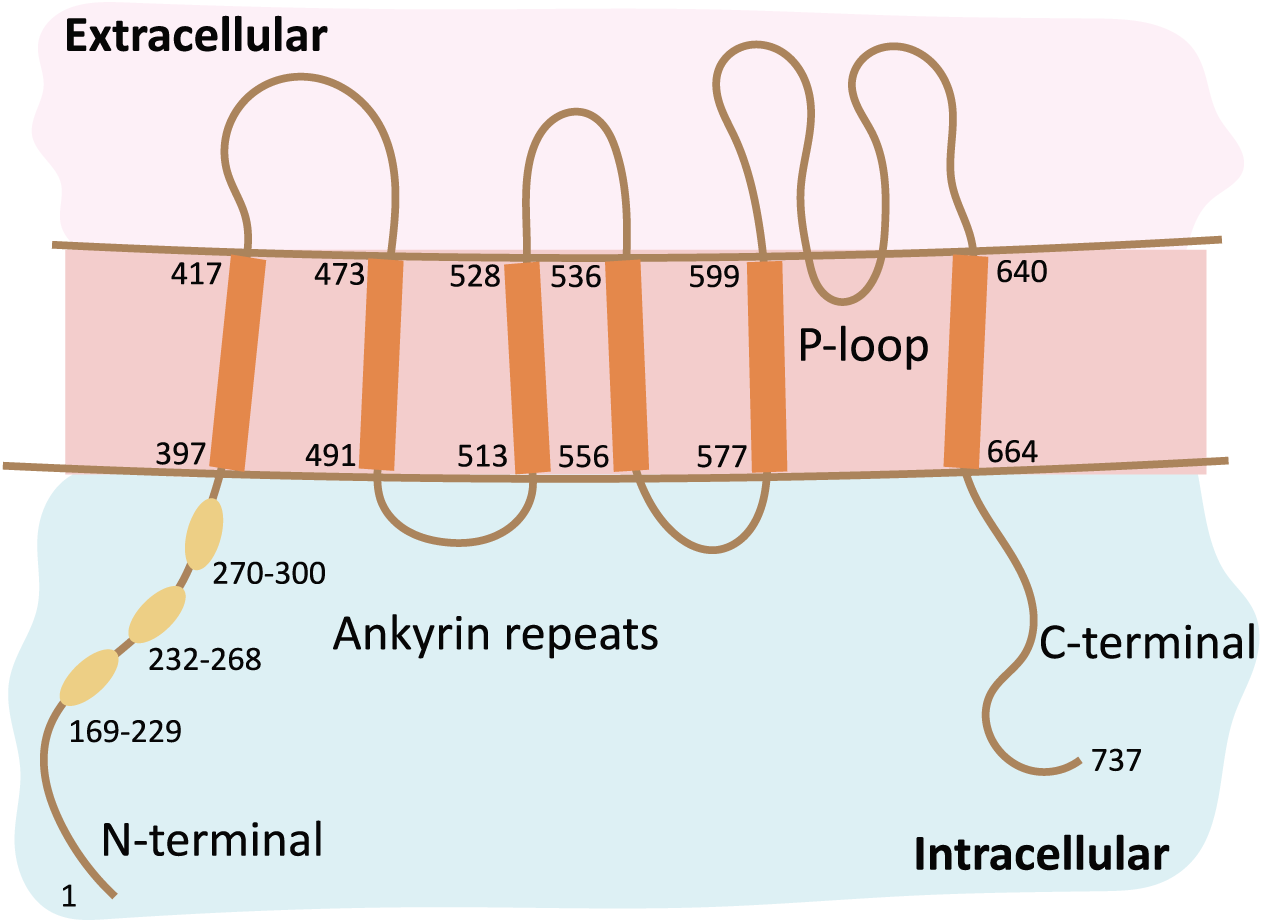
Schematic of the reconstructed *Ovtrpv1* translated protein following secondary and 3D modelling. Typical TRPV channel secondary structure key signatures can be recognised, such as intracellular N- and C-terminals, a variable number of ankyrin repeats in the N-terminal region, six transmembrane elements and a cytosolic P-loop between TM5-TM6 which is responsible for the pore formation and selectivity filter in the assembled tetrameric structure.

### *Ovtrpv1* is located on chromosome 22 of *O. vulgaris* genome

The validated sequence was subsequently blasted against the most recently available *O. vulgaris* genome (Destanović *et al*., 2023) and this led to the identification of a genomic fragment on chromosome 22 (OX597835.1). The translated protein (CAI9738794.1, OctVul6B016571P1) with an automated annotation of “transient receptor potential cation channel subfamily V member 5-like”, was shorter than the sequence predicted and validated. We therefore analysed the putative *Ovtrpv1* transcript against the genomic fragment OX597835.1 using NCBI Magic-BLAST (Boratyn *et al*., 2019). The result confirmed the presence of the transcript distributed in 15 exons (Figure 2A), with an identity of 99.6% over 100% coverage. Only nine mismatches, corresponding to single nucleotide polymorphisms were detected, but these did not affect the translated amino acids. This analysis highlights that the sequence previously tagged by the PV164572 submission and experimentally verified (accession number: PX926345) represents the correct annotation.

**Figure 2.**
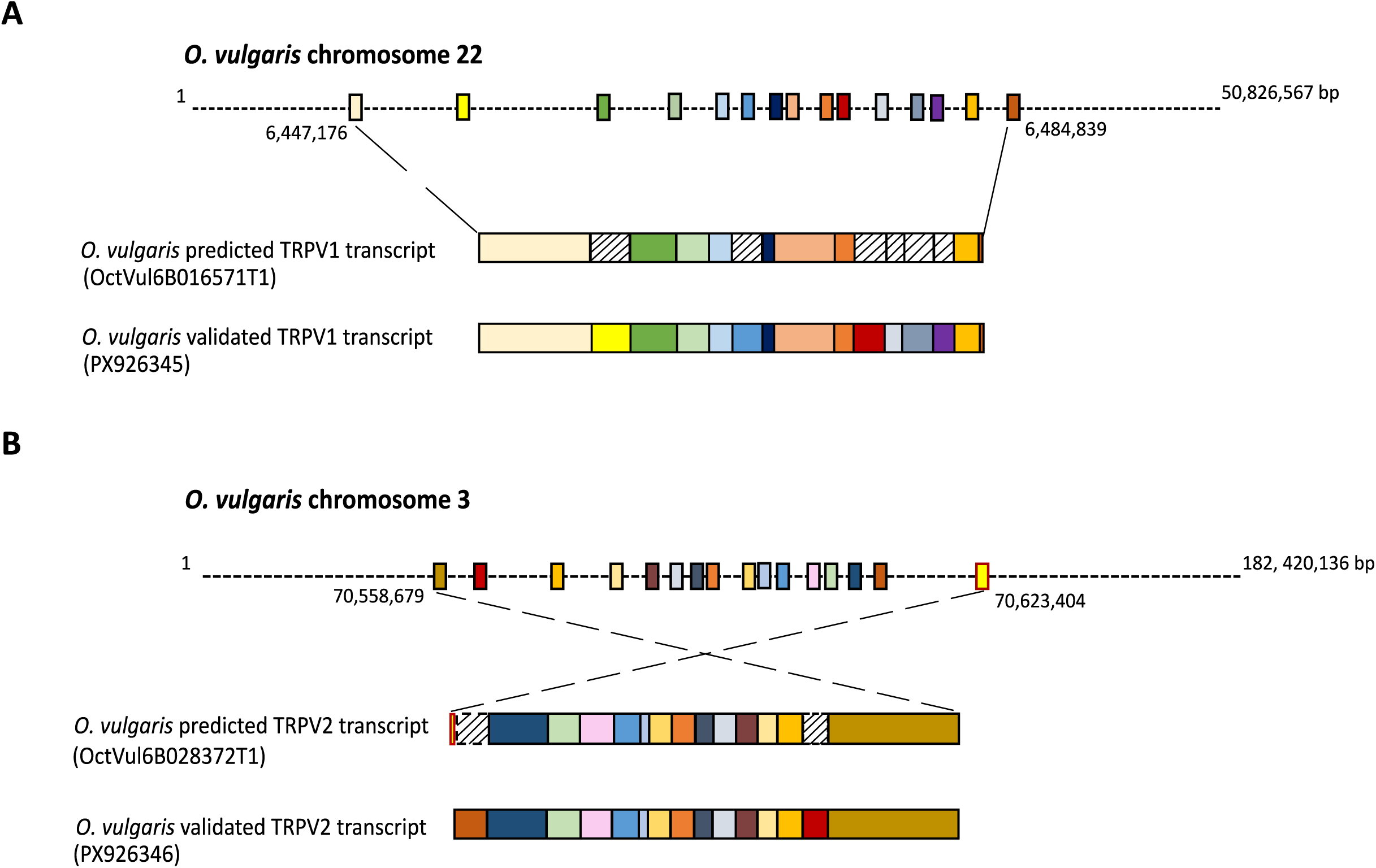
Schematic representation of *Octopus vulgaris trpv1* and *trpv2* gene. **A** *Ovtrpv1* was found to be located on chromosome 22 with 100% coverage and 99.6% identity. **B** The second TRPV candidate gene, *Ovtrpv2*, is located on chromosome 3 and was initially mispredicted (red framed rectangle), requiring experimental validation. Striped pattern rectangles represent initially absent exons.

### *O. vulgaris* genome analysis of *Ovtrpv1* revealed a second TRPV sequence

During our interrogation of the most recent *O. vulgaris* predicted transcriptome (Destanović *et al*., 2023) with *Ovtrpv1*, we identified a distinct sequence located on chromosome 3 (OctVul6B028372T1). This was also automatically annotated based on sequence similarities as “transient receptor potential cation channel subfamily V member 5-like” but was not present in the previously available annotated transcriptome (Petrosino, 2015). Secondary structure prediction and AlphaFold3 modelling utilising the predicted protein sequence, showed again typical TRPV canonical features (Figure 1). However, NCBI conserved domains tool (Lu *et al*., 2020) highlighted homology to the bacterial “Spore_III_AF” super family (cl17562; e-value = 5.81 × 10^-3^) in the predicted protein sequence. To resolve this, we blasted the predicted sequence against other cephalopod transcriptomes and identified a high homology to predicted DNA sequences from *O. sinensis* and *O. bimaculoides*. However, when we aligned the proteins predicted from these DNA sequences, the apparent “Spore_III_AF” super family homology appeared not to be conserved. Therefore, we blasted both *O. vulgaris Ovtrpv* and *O. sinensis* (the closest relative) *Ostrpv* against *O. vulgaris* genome using NCBI Magic-BLAST tool. Both transcripts matched regions of chromosome 3 of *O. vulgaris*. However this matching was not as first envisaged and annotated (Figure 2B). The original *Ovtrpv2* predicted transcript was found to be distributed to 14 exons whilst the subsequent matching listed above found that *Ostrpv2* aligned with an additional exon. Furthermore, exons encoding the N-terminal region of the predicted protein were differently detected, suggesting a potential misprediction of the start codon (Figure 2B). We authenticated this matured prediction using PCR primers designed to flank the ends of the new prediction (Supplementary Table S4). This amplified a PCR product from mRNA derived cDNA of 2799 bp. Sequencing this product confirmed a *bona fide* sequence that does not contain the unexpected bacterial functional domain (Accession number: PX926346). The depiction of the physiological gene structure of *Ovtrpv2* is represented in Figure 2B.

### Cluster analysis and phylogenetic investigation confirmed OvTRPV1 and OvTRPV2 belong to the vanilloid TRPV subfamily

We next asked how these newly identified TRPV sequences related to the other TRP subfamilies. We performed a cluster analysis by blasting canonical curated TRP channel amino acid sequences against the proteome of 13 representative species (Supplementary Table S2). Our CLANS analysis showed the expected distinction among the different subfamilies of TRP channels and from other cationic ion channels (i.e., voltage-gated sodium, calcium and potassium ion channels, Figure 3A). The two newly identified sequences from *O. vulgaris* were found within the same cluster that included other well-characterised vertebrate and invertebrate TRPVs. Our cluster analysis did not identify any additional *O. vulgaris* TRPV candidate. All the sequences belonging to the TRPV cluster, were then used to build a phylogenetic tree to reveal the relationship between the sequences (Figure 3B). The phylogeny suggested invertebrate and vertebrate TRPV receptors share a common ancestor receptor. When looking at the invertebrates, two main branches seemed to have evolved from a duplication event that gave rise to “OCR-like” and “OSM-9-like” TRPV receptor branches, based on the homology with the *C. elegans* TRPV channels. Interestingly, the two OvTRPV sequences were found to belong to distinct branches, with OvTRPV1 belonging to the “OCR-like” and OvTRPV2 belonging to the “OSM-9 like” branch respectively (Figure 3B).

**Figure 3.**
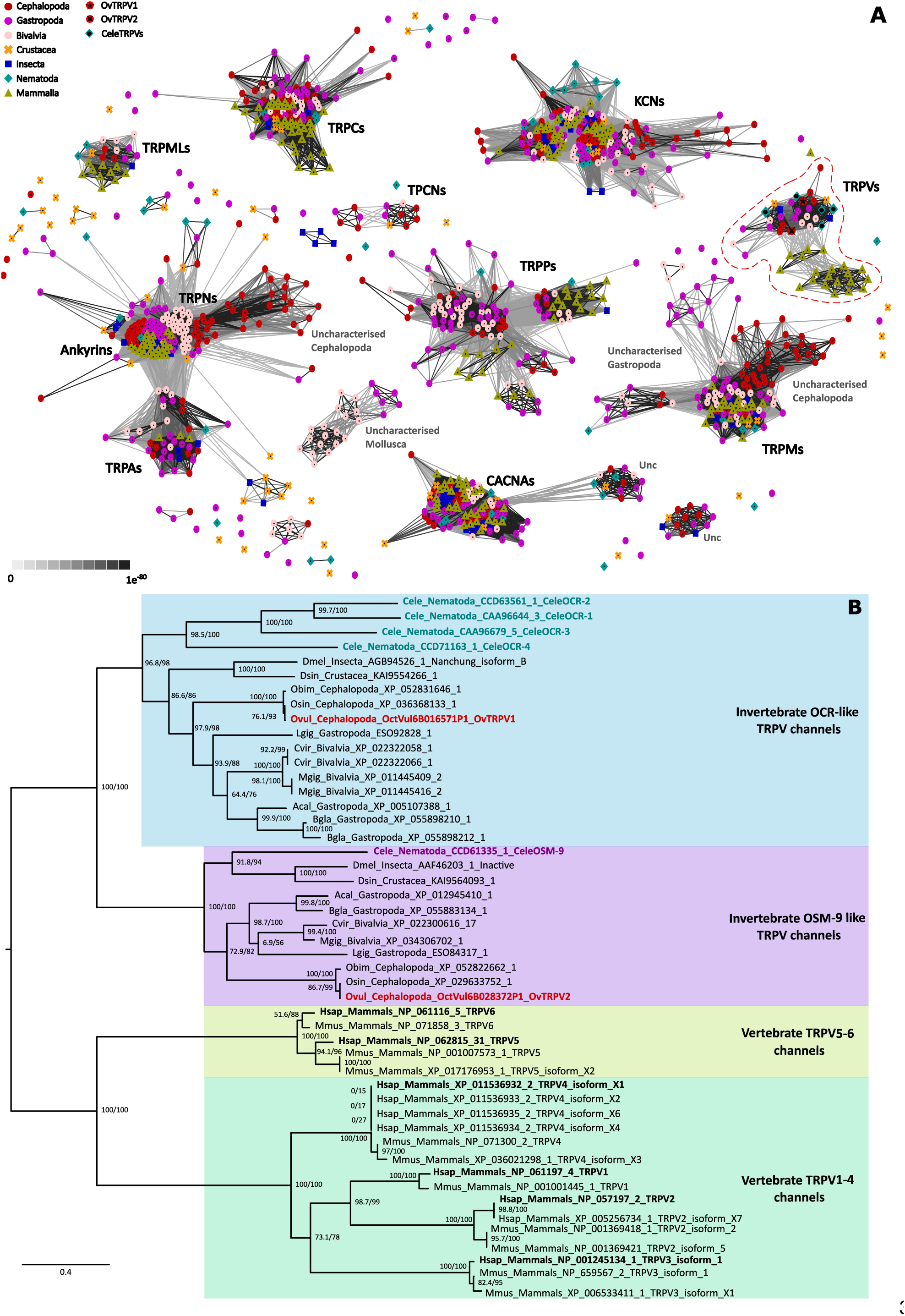
OvTRPV1 and OvTRPV2 are two distinct vanilloid transient receptor potential ion channels. **A** Visual representation of the relationship among the TRP channel subfamilies. The OvTRPVs were identified as part of the TRPV subfamily cluster (dashed red line). The two OvTRPV receptors (red circles with a black star) segregate into two distinct subgroups of the cluster, thus showing some structural distinctions between each other. The same division is observed with the *C. elegans* TRPVs (see legend in the top left corner). CACNA: voltage-gated calcium channels subunit alpha; KCN: potassium voltage-gated channels; TPCN: Two pore calcium channel; TRPA: Transient Receptor Potential cation channel subfamily A; TRPC: transient receptor potential cation channel subfamily C; TRPM: transient receptor potential cation channel subfamily M; TRPML: mucolipin TRP cation channel; TRPP: polycystin transient receptor potential channel interacting; TRPV: transient receptor potential cation channel subfamily V; Unc: uncharacterised. **B** Phylogenetic analysis of OvTRPVs. Phylogenetic tree showing the relationships among TRPV sequences from 13 representative species. The two *O. vulgaris* TRPV sequences (OvTRPV1 and OvTRPV2) are highlighted in bold red. *C. elegans* OCR-like sequences are shown in bold cyan and *C. elegans* OSM-9 in bold purple, while human representative TRPV sequences are indicated in bold black. The tree reveals two distinct TRPV lineages in protostomes, corresponding to “OCR-like” and “OSM-9-like” channels, with OvTRPV1 and OvTRPV2 clustering within these respective groups.

### *Ovtrpv1* and *Ovtrpv2* are expressed in sensory tissues

Once established that the identified sequences are the only readily detected TRPV channel representatives in *O. vulgaris* genome, we investigated their relative tissue distribution using PCR of cDNA synthesised from regionally dissected *O. vulgaris* tissues (Figure 4A). As a comparison, we used an orthologue of the previously identified class of chemotactile receptors (*Ovcrt,* OctVul6B024555T3), found to be selectively expressed in the sensory epithelium of the sucker (van Giesen *et al*., 2020). Our results showed a prevalent expression of both *Ovtrpvs* in the sensory tissues, such as the tip of the arm and the sucker, and in the central brain mass, in a similar fashion to the *Ovcrt* (Figure 4B). Based on the intensity of the genes of interest relative to the control, we evidenced a lower expression in the intestine, white bodies, gill and kidney (Figure 4B).

**Figure 4.**
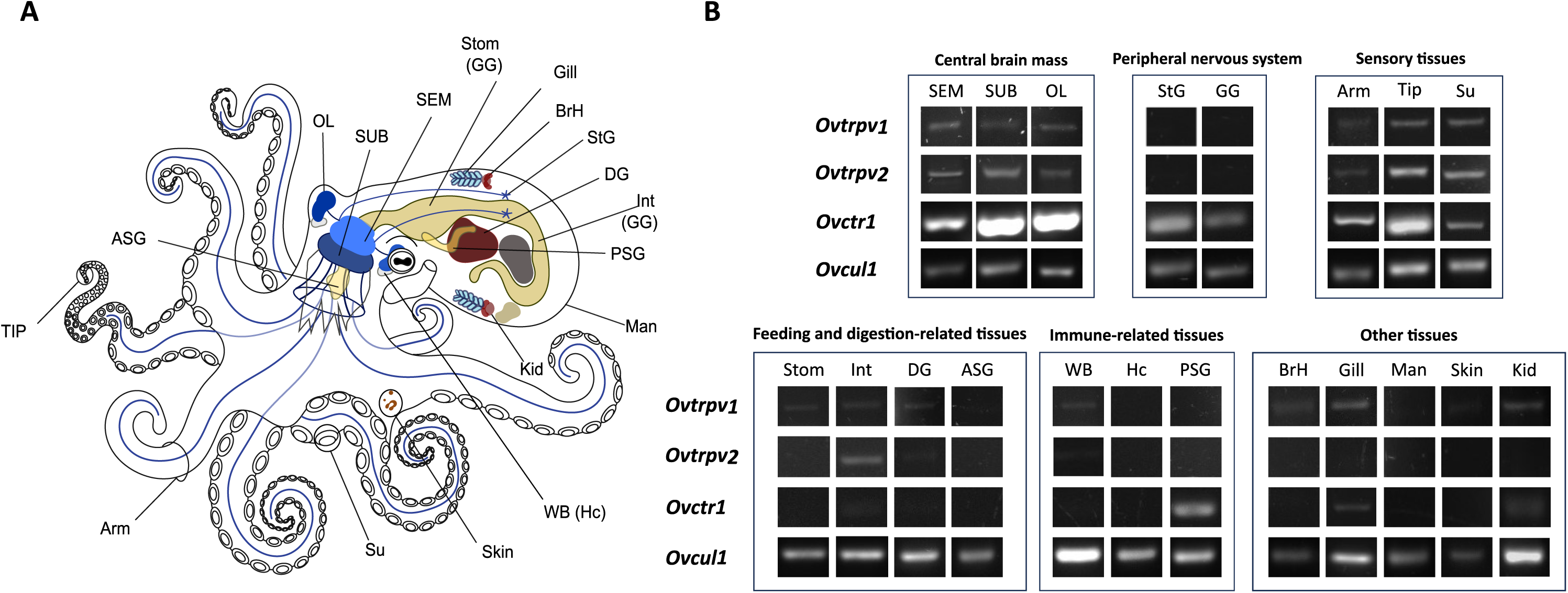
Tissue distribution of *Ovtrpv1* and *Ovtrpv2* receptors in *O. vulgaris.* **A** Schematic representation of *O. vulgaris* tissues localisation in the body. **B** PCR amplification of *Ovtrpv1* and *Ovtrpv2* from a different set of tissue mRNA, grouped in the central brain mass, peripheral nervous system, sensory-, immune- and digestive-related tissues. The data are indicative of the PCR performed on mRNA extracted from the indicated tissue of 4 animals (Supplementary Table S3). Arm: muscle + axial nerve cord, at 50% of its length, TIP: tip of the arm, Su: sucker, OL: optic lobe, SEM: supra-oesophageal mass, SUB: sub-oesophageal mass, StG: stellate ganglion, GG: gastric ganglion, Kid: Kidney, Stom: stomach, Int: intestine, DG: digestive gland, ASG: anterior salivary gland, WB: white bodies, Hc: haemocytes, PSG: posterior salivary gland, BrH: branchial heart, Man: mantle (muscle). *Ovcrt1*: *O. vulgaris* chemotactile receptor 1 (OctVul6B024555T3), *Ovcul1*: *O. vulgaris* cullin 1 (c28856_g1_i2), *Ovtrpv1*: *O. vulgaris trpv1* (PX926345), *Ovtrpv2*: *O. vulgaris trpv2* (PX926346).

### *Ovtrpvs* show polymodal sensory functions when heterologously expressed in *C. elegans*

The tissue distribution described above supports the notion that *Ovtrpv1* and *Ovtrpv2* act as determinants for sensory detection. To investigate this, we took a model hopping approach in *C. elegans*. We previously reported that a functional characterisation of *O. vulgaris* putative nociceptive genes can be carried out in *C. elegans* using behavioural readouts (Pieroni *et al*., 2026). We found high sequence homology to the nociceptive-related TRPV channels *Celeocr-2* and *Celeosm-9*. OvTRPV1 shares 55.0% amino acid similarity and 39.8% amino acid identity with the sensory OCR receptor CeleOCR-2, while OvTRPV2 shares 54.7% amino acid similarity, and 40.5% amino acid identity with CeleOSM-9. For this reason, we heterologously expressed *Ovtrpv1* and *Ovtrpv2* cDNA sequence in the corresponding *C. elegans ocr-2 (ak47)* and *osm-9 (ky10)* loss of function mutants under the control of their putative respective worm promoters and tested the resulting lines for rescue of avoidance to different nociceptive modalities.

We first investigated the ability of *Ovtrpv* expression to restore avoidance to low pH, a cue previously identified as relevant to cephalopod nociception (Hague, Florini and Andrews, 2013; Crook, 2021). Aversion to low pH was measured using a classical acute aversion assay, namely the drop assay. WT N2 worms responded around 70% of the time by displaying 3 or more backward movements once in contact with the drop of nociceptive cue (Figure 5A and B). The *ocr-2* (A) and *osm-9* (B) mutants showed reduced backing response when the worms were exposed to pH 3, consistent with previous data (Sambongi *et al*., 2000). This was also observed in the mutant strains carrying the *gfp* transgene which were used as a reference for the rescue (*ocr-2 (ak47)* vs *ocr-2 (ak47) Pmyo-3::gfp*, p= 0.9773; *osm-9 (ky10)* vs *osm-9 (ky10) Pmyo-3::gfp*, p= 0.9878). When we reintroduced the *C. elegans* gene as a fosmid (*ocr-2 (WRM0634bB10) Pmyo-3::gfp* vs *ocr-2 (ak47) Pmyo-3::gfp,* p<0.0001*; osm-9 (WRM066bG12) Pmyo-2::gfp* vs *osm-9 (ky10) Pmyo-3::gfp*, p<0.0001) or cDNA (*Pocr-2::Celeocr-2 Pmyo-3::gfp* vs *ocr-2 (ak47) Pmyo-3::gfp*, p<0.0001; *Posm-9::Celeosm-9 Pmyo-3::gfp* vs *osm-9 (ky10) Pmyo-3::gfp,* p= 0.0006) under the respective *ocr-2* and *osm-9* promoter, we restored the reduced pH sensitivity (Figure 5A and B).

**Figure 5.**
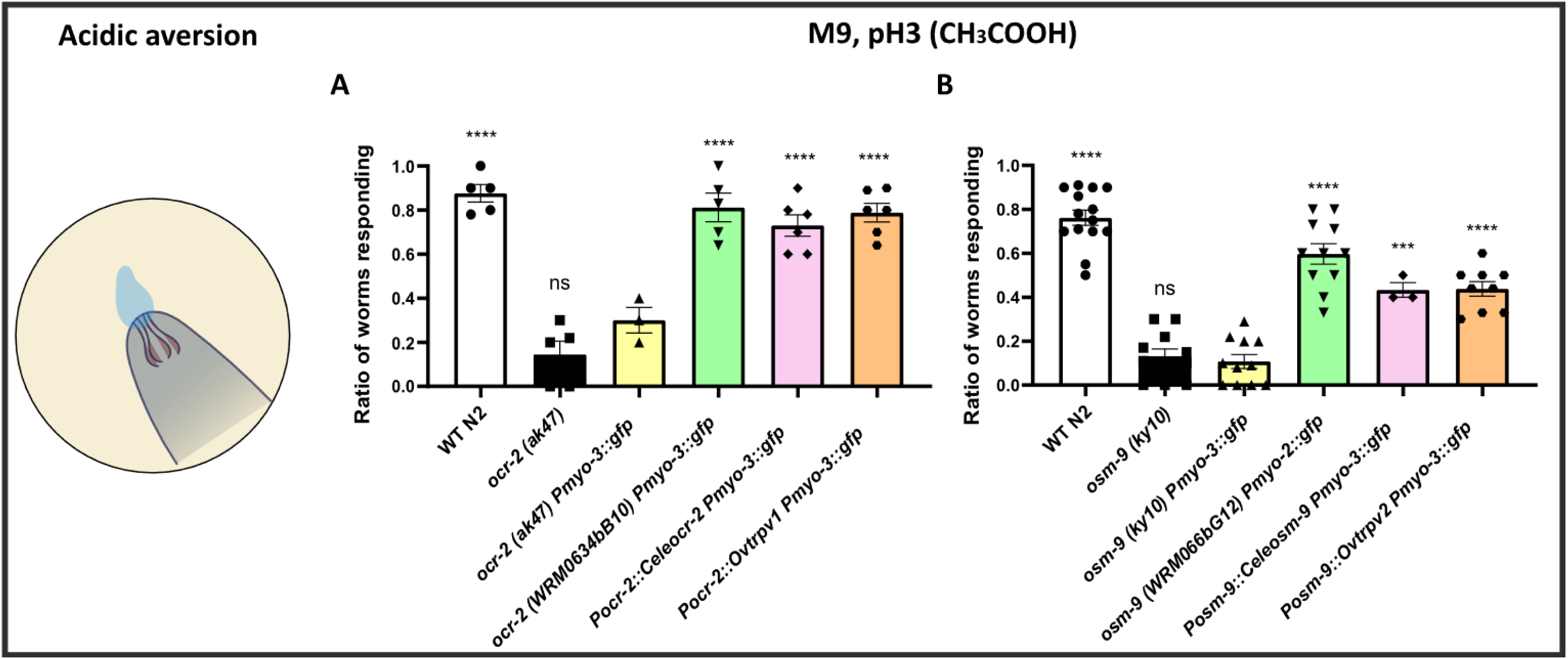
*Ovtrpv1* and *Ovtrpv2* rescue pH3-evoked avoidance in *C. elegans ocr-2* (A) and *osm-9* (B) mutant strains. The bar graphs represent the ratio of worms responding ± s.e.m. for WT N2*, ocr-2 (ak47), osm-9 (ky10), ocr-2 (ak47) Pmyo-3::gfp, osm-9 (ky10) Pmyo-3::gfp* and fosmid lines. Each dot represents a replicate experiment in which we tested 10 worms per condition. For the *C. elegans* and *O. vulgaris* cDNA constructs, each dot represents a rescue line (10 worms each), which was selected according to the threshold set at 2 standard deviations above the mean of the reference mutant line *ocr-2 (ak47) Pmyo-3::gfp* (A) or *osm-9 (ky10) Pmyo-3::gfp* (B). All the average performances are here compared to the reference mutant lines. Data were analysed using one-way ANOVA and Post-hoc comparisons have been performed with Dunnett’s multiple comparisons test. ***p<0.001, **** p<0.0001.

Against this background we repeated the analysis with cDNA encoding OvTRPV1 and OvTRPV2. Worm lines containing the *Ovtrpvs* cDNA showed a significant rescue of the response in the mutant lines (*Pocr-2::Ovtrpv1 Pmyo-3::gfp vs ocr-2 (ak47) Pmyo-3::gfp,* p<0.0001*; Posm-9::Ovtrpv2 Pmyo-3::gfp vs osm-9 (ky10) Pmyo-3::gfp,* p<0.0001).

In detail, *Ovtrpv1* led to a full rescue of the phenotype (*Pocr-2::Ovtrpv1 Pmyo-3::gfp vs WT N2* p= 0.6235, Figure 5A).*Ovtrpv2* showed a significant difference to the WT performance (*Posm-9::Ovtrpv2 Pmyo-3::gfp vs osm-9 (ky10) Pmyo-3::gfp,* p<0.0001) suggesting a clear but incomplete rescue. However, this was also true for the corresponding *C. elegans* cDNA (*Posm-9::Celeosm-9 Pmyo-3::gfp* vs WT N2, p= 0.0004) and genomic (*osm-9 (WRM066bG12) Pmyo-2::gfp* vs WT N2, p= 0.0068) rescue constructs (Figure 5B). Thus, the partial rescue may reflect methodological confounds of the transgenic expression (e.g., mosaic expression of the transgene, missing additional regulatory elements in the promoter region), rather than a reduced ability of the orthologue to substitute function.

Next, mechanical aversion was investigated in *C. elegans* using a nose touch assay, another classical acute aversion test (Figure 6). Similarly to low pH aversion, WT worms showed an halted and/or backing behaviour when in contact with the eyebrow (Figure 6A and B).

**Figure 6.**
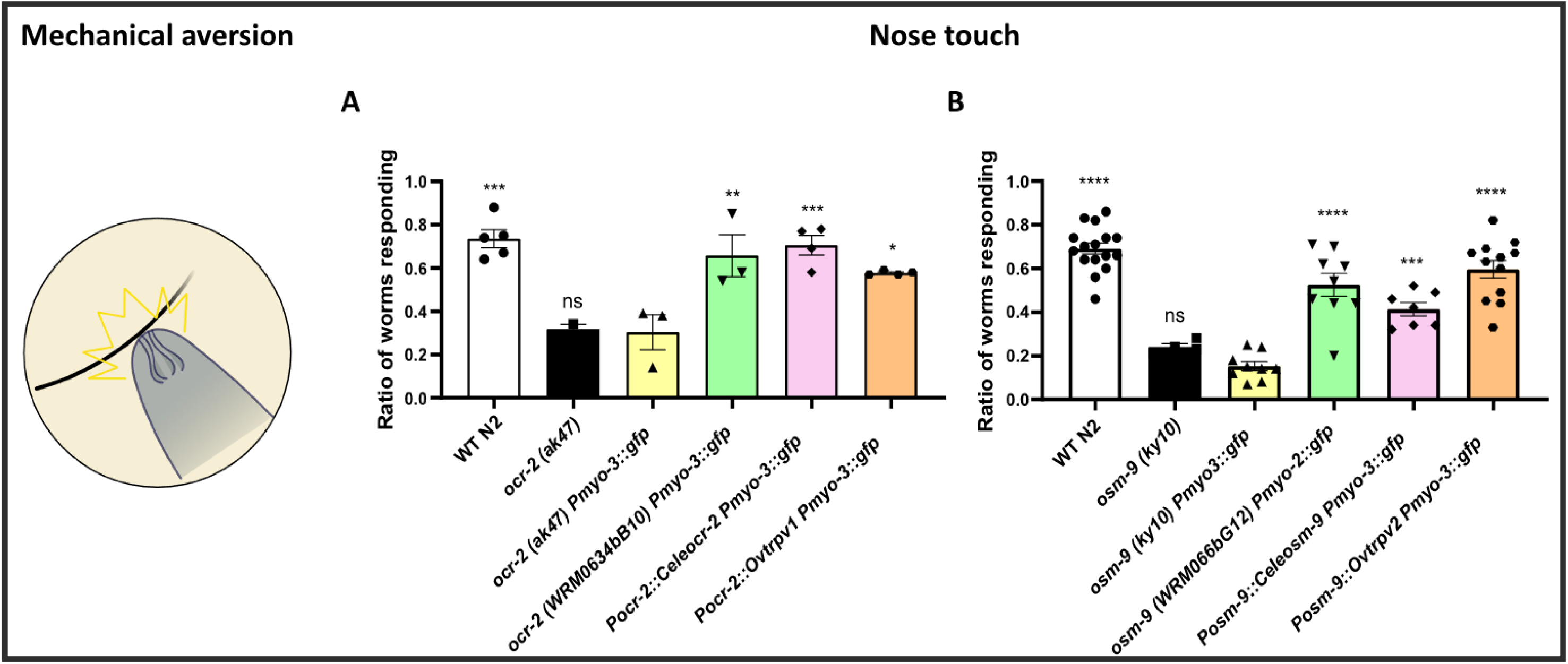
*Ovtrpv1* and *Ovtrpv2* rescue mechanical aversion in *C. elegans ocr-2* (A) and *osm-9* (B) mutant strains. The bar graphs represent the ratio of worms responding ± s.e.m. for WT N2*, ocr-2 (ak47), osm-9 (ky10), ocr-2 (ak47) Pmyo-3::gfp, osm-9 (ky10) Pmyo-3::gfp*, fosmid lines and *Ovtrpv* expressing lines. Each dot represents a replicate experiment in which we tested 5 worms in a set of 10 trials. For the *C. elegans* and *O. vulgaris* cDNA constructs, each dot represents a rescue line (5 worms per 10 trial each), which was selected according to the threshold set at 2 standard deviations above the mean of the reference mutant line *ocr-2 (ak47) Pmyo-3::gfp* (A) or *osm-9 (ky10) Pmyo-3::gfp* (B). All the average performances are here compared to the reference mutant lines. Data were analysed using one-way ANOVA and Post-hoc comparisons have been performed with Dunnett’s multiple comparisons test *p<0.05, **p<0.01, ***p<0.001, **** p<0.0001.

Again, both mutant lines showed an impaired response to the mechanical insult (*ocr-2 (ak47)* vs *ocr-2 (ak47) Pmyo-3::gfp*, p>0.9999; *osm-9 (ky10)* vs *osm-9 (ky10) Pmyo-3::gfp*, p= 0.5753) a defective response already reported for both mutant strains (Tobin *et al*., 2002).

The introduction of the fosmid for both *ocr-2* and *osm-9* genes (Figure 6A and B) was able to recover the lost aversive response in both mutant strains (*ocr-2 (WRM0634bB10) Pmyo-3::gfp* vs *ocr-2 (ak47) Pmyo-3::gfp,* p<0.0032*; osm-9 (WRM066bG12) Pmyo-2::gfp* vs *osm-9 (ky10) Pmyo-3::gfp*, p<0.0001). Similarly, *C. elegans ocr-2* and *osm-9* cDNA showed a successful rescue (*Pocr-2::Celeocr-2 Pmyo-3::gfp* vs *ocr-2 (ak47) Pmyo-3::gfp*, p= 0.0005; *Posm-9::Celeosm-9 Pmyo-3::gfp* vs *osm-9 (ky10) Pmyo-3::gfp,* p= 0.0002).

When introducing the *O. vulgaris trpv* cDNA, both constructs were able to rescue the gene function by recovering aversion to mechanical insults. *Ovtrpv1* exhibited a modest rescue in *ocr-2 (ak47)* background (*Pocr-2::Ovtrpv1 Pmyo-3::gfp vs ocr-2 (ak47) Pmyo-3::gfp,* p= 0.0131, Figure 6A) while *Ovtrpv2* showed a full rescue of the behavioural phenotype of *osm-9 (ky10)* with more than 70% of the lines tested (12 out of 17) efficiently responding to nose touch (*Posm-9::Ovtrpv2 Pmyo-3::gfp vs WT N2,* p= 0.1497, Figure 6B).

We additionally tested rescue to another chemical noxious compound, the volatile repulsive cue 1-octanol by measuring the latency to start a reversal behaviour when exposed to the airborne compound (Figure 7). The impairment of the mutant strains was striking (average latency of *ocr-2 (ak47) Pmyo-3::gfp* of 11.65 s vs average latency of 4.13 s of WT N2; average latency of *osm-9 (ky10) Pmyo-3::gfp of* 11.06 s vs average latency of 2.25 s of WT N2, Figure 7A and B) and has been previously reported in *C. elegans* TRPV mutants (Thies *et al*., 2016).

**Figure 7.**
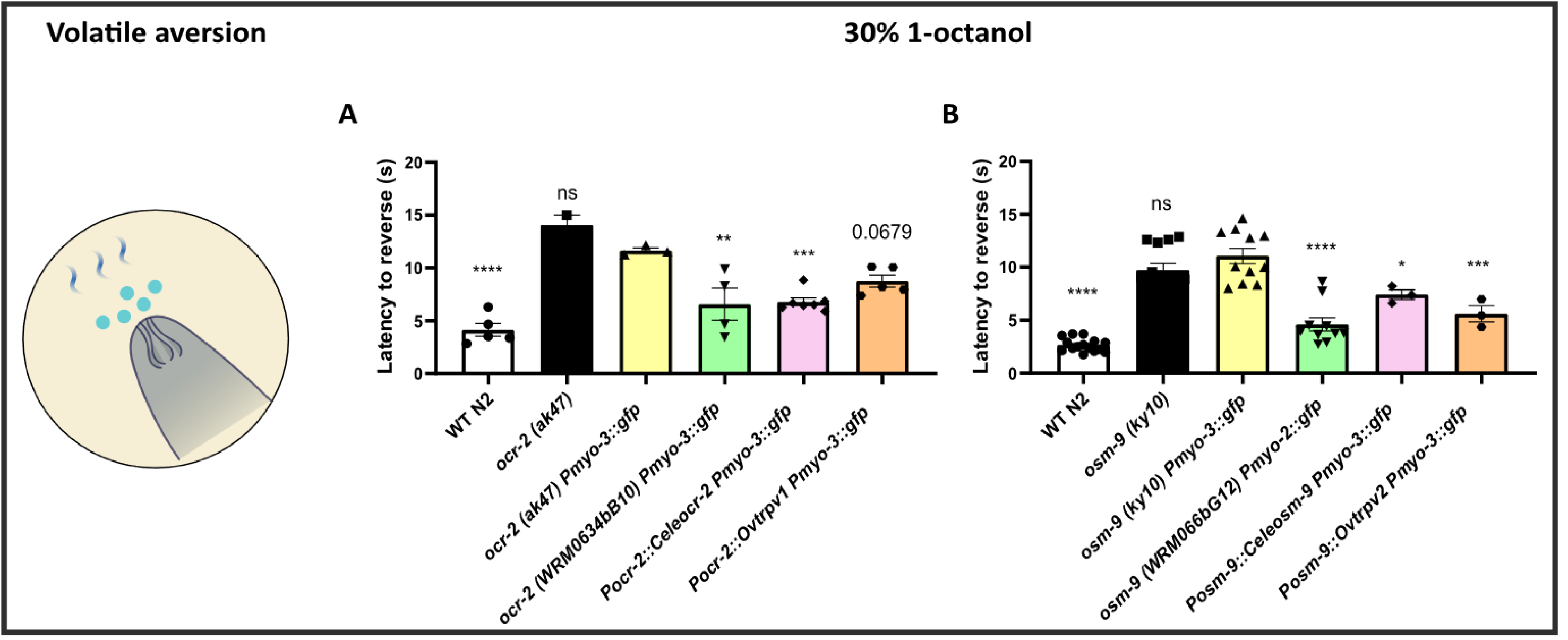
Investigation *of* volatile aversion in *C. elegans ocr-2 (A)* and *osm-9* (B) mutant strains. The bar graphs represent the average latency to start a reversal ± s.e.m. for WT N2*, ocr-2 (ak47), osm-9 (ky10), ocr-2 (ak47) Pmyo-3::gfp, osm-9 (ky10) Pmyo-3::gfp*, fosmid lines and *Ovtrpv* expressing lines. Each dot represents a replicate experiment in which we tested 5 worms in a set of 10 trials. For the *C. elegans* and *O. vulgaris* cDNA constructs, each dot represents a rescue line (10 worms each), which was selected according to the threshold set at 2 standard deviations below the mean of the reference mutant line *ocr-2 (ak47) Pmyo-3::gfp* (A) or *osm-9 (ky10) Pmyo-3::gfp* (B). All the average performances are here compared to the reference mutant lines. Data were analysed using one-way ANOVA and Post-hoc comparisons have been performed with Dunnett’s multiple comparisons test *p<0.05, **p<0.01, ***p<0.001, **** p<0.0001.

Rescue was achieved with both genomic (*ocr-2 (WRM0634bB10) Pmyo-3::gfp* vs *ocr-2 (ak47) Pmyo-3::gfp,* p= 0.0016*; osm-9 (WRM066bG12) Pmyo-2::gfp* vs *osm-9 (ky10) Pmyo-3::gfp*, p<0.0001) and cDNA construct (*Pocr-2::Celeocr-2 Pmyo-3::gfp* vs *ocr-2 (ak47) Pmyo-3::gfp*, p<0.0009; *Posm-9::Celeosm-9 Pmyo-3::gfp* vs *osm-9 (ky10) Pmyo-3::gfp,* p= 0.0172).

When heterologously expressing octopus cDNA, only a trend towards a reduced latency to initiate reversals to diluted (30%) 1-octanol was shown by *Ovtrpv1*-expressing lines (*Pocr-2::Ovtrpv1 Pmyo-3::gfp vs ocr-2 (ak47) Pmyo-3::gfp,* p= 0.0679, Figure 7A), while *osm-9 (ky10)* mutants expressing *Ovtrpv2* were successfully rescued (*Posm-9::Ovtrpv2 Pmyo-3::gfp vs osm-9 (ky10) Pmyo-3::gfp*, p= 0.0002, Figure 7B).

### Investigation of OvTRPVs function using recombinant systems

The successful rescue of different modalities and cues through expression of *Ovtrpv* channels in mutant *C. elegans* strains, suggested they share sufficient structural similarity with their worm orthologues to function in a similar fashion to the missing endogenous receptors in the intact organism. However, whether these act upstream as direct activators or downstream as final effectors or modulators of the primary sensory signalling is not known. To address this, we heterologously expressed the receptors in recombinant systems, as it was previously shown in *C. elegans* that OSM-9/OCR-2 are both required to be correctly localised at the ciliated sensory neurons to exert a functional response (Tobin *et al*., 2002; Ohnishi *et al*., 2020). We used *Xenopus* oocytes to test for direct receptor activation. We verified the ability to reconstitute a ligand-activated response from the *Homo sapiens Hstrpv1* control using capsaicin, but *Ovtrpvs*-expressing oocytes did not respond to this vanilloid compound, suggesting this might not be an ecologically relevant stimulus in cephalopods (Figure 8A). However, a long-sustained response to 100 μM nicotinamide (NAM) was observed when *Ovtrpv1* and *Ovtrpv2* were co-expressed but not when either receptor was expressed alone (Figure 8B), suggesting that OvTRPV subunits require co-assembly of the two subunits to reconstitute a response in a recombinant system.

**Figure 8.**
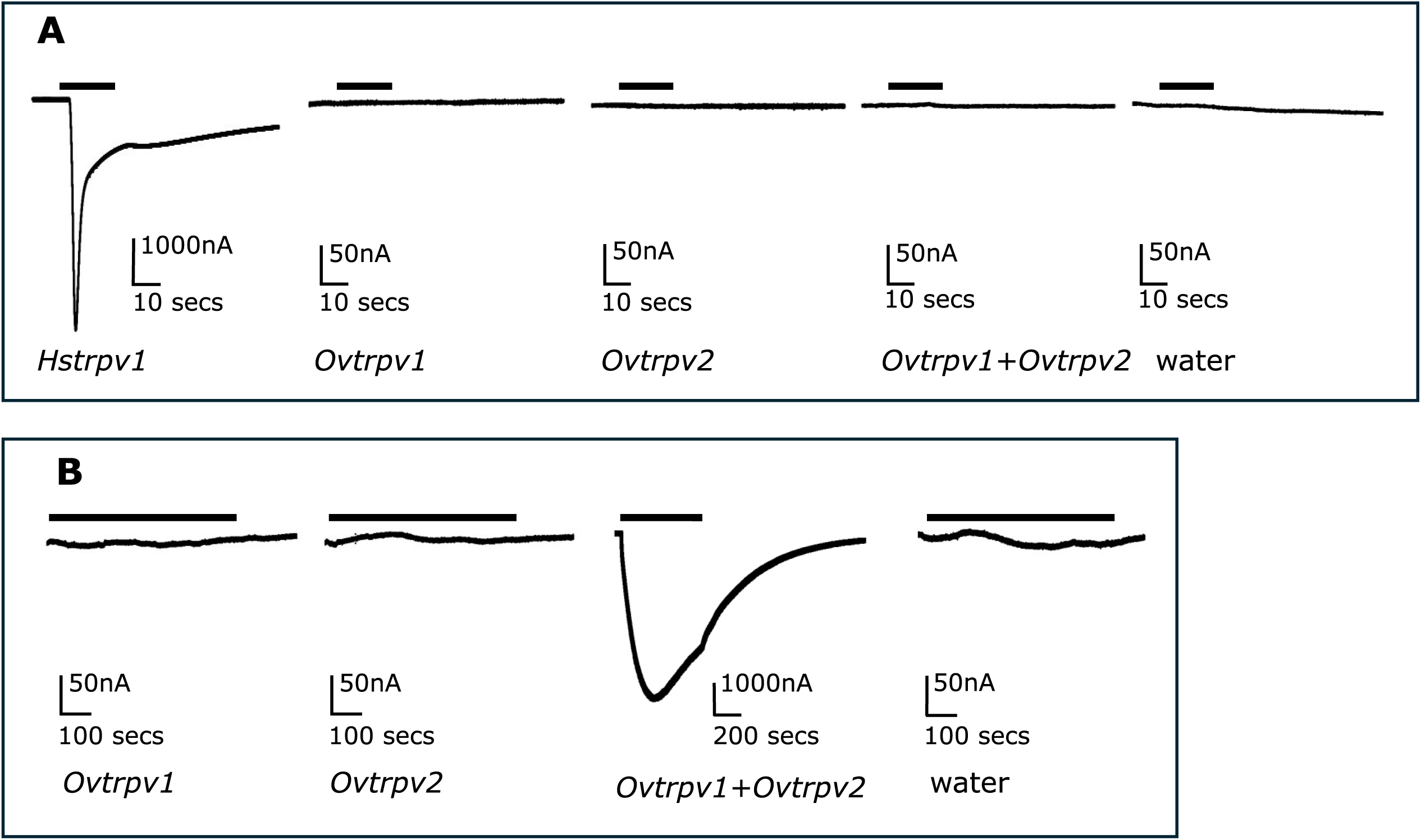
Heterologous expression of *O. vulgaris* TRPV channels in *Xenopus* oocytes. **A.** Current traces from *Xenopus* oocytes injected with RNA encoding the human TRPV1 channel shows response to capsaicin at 100µM. The solid bars represent the period of capsaicin perfusion. Oocytes injected with RNA for *O. vulgaris trpv1* and *trpv2* separately or together show no response to capsaicin at 100µM. Oocytes injected with water show no response. N ≥ 7 oocytes for each condition. **B.** *O. vulgaris* TRPV1 and TRPV2 form a heteromeric channel which responds to nicotinamide. *Xenopus* oocytes injected with RNA for *Ovtrpv1*, *Ovtrpv2* or water show no responses to nicotinamide at 100µM. Oocytes expressing *Ovtrpv1* and *Ovtrpv2* together show strong responses to nicotinamide 100µM. N ≥ 7 oocytes for each condition.

## Discussion

### *O. vulgaris* has two distinct TRPV channel representatives

In a previous *in silico* analysis of the conserved molecular determinants putatively involved in *O. vulgaris* nociception, we highlighted candidates that could underpin octopus response to noxious stimuli (Pieroni *et al*., 2026).

In this study, we focussed on the nociceptive molecular determinant TRPV that belongs to the vanilloid subfamily of TRP channels (Himmel and Cox, 2020).

The TRPV subfamily, and particularly the TRPV1 member in vertebrates, is known to be gated by several nociceptive cues including capsaicin, low pH and high temperature, thus classifying it as a polymodal receptor (Caterina *et al*., 1997; Immke and Gavva, 2006; Dhaka *et al*., 2009). An important aspect of TRPV1 and related proteins is that this molecular sensing and signalling is placed in discrete cells in the peripheral part of the body so they can act as sensory receptors that initiate complex downstream signalling that allows the embedded circuits to drive judicious behavioural responses (Caterina *et al*., 1997). Consistent with this, a prevalent expression of TRPV1 in peripheral sensory neurons that interact with the environment has been extensively reported (Caterina *et al*., 1997; Nakagawa and Hiura, 2006; Szigeti *et al*., 2012). In addition, a wider expression in the central brain implying distinct contributions to nociception or other functions has also been found (Yuan and Burrell, 2010; Higgins *et al*., 2013; Marrone *et al*., 2017).

The sensory function and the anatomical distribution of TRPV channels also pertain to invertebrate and simpler organisms (Colbert, Smith and Bargmann, 1997; Summers, Holec and Burrell, 2014). This prompted us to investigate the TRPV subfamily in *O. vulgaris* in which its complex nervous system and articulated behaviours are speculated to allow elaboration of nociception into pain-like states (Birch *et al*., 2021). Our work exploited the emerging access to octopus transcriptome and genome databases and matured earlier designation of a single TRPV gene. Cluster analysis and phylogenetic investigation refined the designation of OvTRPV1 and identified a second receptor, OvTRPV2. Our analysis authenticated these genes and highlighted they are the only representatives of vanilloid TRP channels found in *O. vulgaris* and represent two discrete receptor channel subunits (Figure 3A). Two TRPV representatives were also found in other cephalopods such as *O. bimaculoides* and *O. sinensis* and also in other species such as *D. melanogaster* or *A. californica* (Figure 3B). This indicates potential structural conservation across distinct invertebrate genera which may well reflect strong functional conservation of sensory specialisation.

An interesting exception to the generalisation above is represented by *C. elegans*, which shows a total of 5 TRPV representatives, suggesting a specific one-to-many orthologues expansion of these receptors compared to other invertebrate phyla (Kuzniar *et al*., 2008). We can identify an invertebrate TRPV group referred to as “OCR-like” TRPV branch. This grouping includes all the *C. elegans* OCR receptors (OCR 1-4) to which OvTRPV1 and *D. melanogaster* Nanchung (NAN) are more closely related (55.0% similarity). A second group, referred to as “OSM-9 like” branch, includes *C. elegans* OSM-9 to which OvTRPV2 and *D. melanogaster* Inactive (IAV) are related (54.7% similarity, Figure 3B). The analysis shows the invertebrate TRPVs share a common ancestor with the vertebrate TRPVs. This analysis supports a scenario in which this ancestor receptor underwent a duplication that gave rise to two distinct mammalian branches. One led to the TRPV5 and 6 ion channels which, despite retaining sensitivity to pH, do not have any sensory related functions but act to regulate calcium homeostasis in the kidney and small intestine (Yeh *et al*., 2003; Nijenhuis, Hoenderop and Bindels, 2005; Lambers *et al*., 2007). This subgroup is separated from the TRPV1-4 channels branch, in which further internal duplications likely led to a specialisation of these receptors in compounds and stimuli detection (Peng, Shi and Kadowaki, 2015; Morini *et al*., 2022).

### Tissue expression supports the hypothesis that *Ovtrpvs* are sensory molecular determinants in octopus

As indicated, the tissue expression of the mammalian TRPV1 justifies key function in sensory signalling (Caterina *et al*., 1997). In the case of the *Ovtrpvs*, gross analysis of their mRNA expression across different tissues supports an important role in sensory signalling (Figure 4). Both subunits appear to show relatively low expression but robust and reproducible amplification from tissues associated with sensory signalling in cephalopods. This is reinforced when we compare the expression in the sensory tissues (arm, tip of the arm and sucker) to a range of tissues encapsulating immune-related (e.g., white bodies, haemocytes) and feeding and digestion-related (e.g., stomach, intestine) tissues (Figure 4B). Although tissue distribution cannot define specificity of function *per se*, we identified that *Ovtrpv1* and *Ovtrpv2* have overlapping expression with the selective chemotactile receptor orthologue *Ovcrt* (Figure 4B). This gene is reported to be enriched in the sensory epithelium of the sucker of *O. bimaculoides* with reported related sensory functions (van Giesen *et al*., 2020). This enrichment is interesting as cephalopods, and in particular octopuses, use their arms to sample the environment and classify stimuli including those that evoke aversion. These structures which express sensory receptors are primary organisers of reflexive and complex responses (Rossi and Graziadei, 1956; Zullo, Fossati and Benfenati, 2011; Hague, Florini and Andrews, 2013; Gutnick *et al*., 2020). The prevalence of *Ovtrpv1* and *Ovtrpv2* expression in the distal part of the arm, such as the tip and more specifically the suckers, encourages the hypothesis these receptors modulate sensory detection in *O. vulgaris*. The comparison with *Ovcrt* expression suggests these receptors are low abundant, and this seems in line, at least for *Ovtrpv1,* with the RNA sequencing analysis of its fragments performed in the previously assembled *O. vulgaris* transcriptome (Petrosino, 2015; Petrosino *et al*., 2022).

In a similar fashion to vertebrate TRPVs we showed that, *Ovtrpvs* expressed in the central brain mass of *O. vulgaris* (Figure 4B), supports a potential role in central control of sensory and nociceptive modulation.

We additionally detected expression of both *Ovtrpvs* in the intestine and, exclusively for *Ovtrpv1*, in the stomach, digestive gland, white bodies, gill and kidney (Figure 4B). This broader expression in non-neuronal tissues for *Ovtrpv1* is interesting when considering it belongs to the “OCR-like” branch. In *C. elegans*, OCRs show a more heterogenous expression including the rectal gland cells and intestine which play an important role in *C. elegans* digestive system (Tobin *et al*., 2002; Packer *et al*., 2019). Additionally, *ocr-3* and *ocr-4* seem to be co-expressed in neuroendocrine cells with *ocr-2,* to regulate egg-laying (Jose *et al*., 2007).

### Functional characterisation through model hopping in *C. elegans* suggests

#### *Ovtrpvs* are polymodal nociceptors

The prevalent expression in sensory tissues of both receptors, encouraged us to functionally test them for their role in nociception.

As we have already utilised *C. elegans* as a suitable model to perform functional characterisation of octopus molecular nociceptor candidates (Pieroni *et al*., 2026), we resorted to a model hopping approach by expressing octopus receptors in the mutant strains for the orthologue genes *ocr-2* and *osm-9*. The phylogenetic distinction between the two receptors into “OCR-like” receptors for *Ovtrpv1* and “OSM-9-like” for *Ovtrpv2*, justified our approach of heterologously expressing *Ovtrpv1* under the *ocr-2* promoter in an *ocr-2 (ak47)* mutant background and *Ovtrpv2* under the *osm-9* promoter in an *osm-9 (ky10)* mutant background.

The null mutants *osm-9 (ky10)* and *ocr-2 (ak47)* show defects in different sensory modalities, including chemical and mechanical aversion, effectively highlighting OSM-9 and OCR-2 as polymodal receptors (Colbert, Smith and Bargmann, 1997; Tobin *et al*., 2002; Liedtke *et al*., 2003; Thies *et al*., 2016). We therefore subjected our transgenic lines expressing *Ovtrpv1* and *Ovtrpv2* in the *C. elegans* mutants for the orthologous genes to acetic acid (pH 3), mechanical insults (i.e. nose touch) and volatile repellents such as 1-octanol.

*Ovtrpv1* and *Ovtrpv2* successfully rescued low pH (Figure 5) and mechanical avoidance (Figure 6) suggesting these receptors act as polymodal nociceptors in *O. vulgaris*. These two cues induce aversive responses in cephalopods. Low pH has been tested in *ex vivo* octopus arm preparations, showing a quick withdrawal when in contact with an acidic solution, and *in vivo* experiments, in which injection of acetic acid triggered grooming and protective behaviours (Hague, Florini and Andrews, 2013; Crook, 2021). Mechanical cues are also an important trigger for cephalopod sensory responses as von Frey filaments induce nociceptive responses and also trigger post-injury sensitisation in squids (Crook, Hanlon and Walters, 2013; Alupay, Hadjisolomou and Crook, 2014; Crook *et al*., 2014).

However, the molecular determinants of such responses have not been investigated in *O. vulgaris* and the TRPV channels we describe here are potential candidates. Whether OvTRPVs act as channels directly gated by pH or as indirect modulators of pH behavioural responses is not known and the proposed key residues for proton gating in mammalian TRPV1 (Jordt and Julius, 2002; Ryu *et al*., 2007; Aneiros *et al*., 2011), are not conserved in OvTRPVs or in CeleTRPVs. Therefore, either OvTRPV channels have a distinct mechanism for pH detection, or perhaps they possess a downstream modulatory role for this cue. As for mechanosensation, a key molecular determinant has been found in *O. bimaculoides* as part of the TRPN family, which also includes the mechanosensory *D. melanogaster* NompC (Yan *et al*., 2013; van Giesen *et al*., 2020). However, it is not uncommon in organisms to have multiple receptors responding to the same cue. This allows resolution of the stimulus detection and its texture, that leads to the distinction between a soft touch and a nociceptive harmful input. *C. elegans* represents a good example of this, as *osm-9/ocr-2* are not uniquely required for aversive nose touch (Kindt *et al*., 2007; Li *et al*., 2011), and different molecular interactions give rise to distinct sensitivities (e.g., *mec-10/mec-4* for gentle touch and *mec-10/degt-1* for harsh touch, Chatzigeorgiou *et al*., 2010; Li *et al*., 2011).

Our *C. elegans* model hopping suggests that homologue expression gives the most potent rescue with *ocr-2* promoter driving *Ovtrpv1* and *osm-9* promoter driving *Ovtrpv2* expressions in nematode loss of function mutants of the orthologue *trpv* genes. However, these functional experiments preclude insight into their transduction localisation (upstream or downstream) or the required stoichiometry that makes the receptor functional in sensory cellular signalling. The evolutionary relationship between *Ovtrpv1* and *Ovtrpv2,* consistent with other invertebrates, suggests important divergence in function (Figure 3B). Furthermore, previous experiments suggested co-assembly of *osm-9* and *ocr-2* to exert their role (Ohnishi *et al*., 2020; Griffin *et al*., 2025). Finally, the shared overlapping tissue expression of *Ovtrpv1* and *Ovtrpv2* suggests they might interact together to convey sensory detection. Altogether, these pieces of evidence, encouraged our approach of using recombinant assays.

Heterologous co-expression of *Ovtrpv1* and *Ovtrpv2* in *Xenopus* oocytes produced a sustained, slow response to 100 μM NAM, supporting the hypothesis of co-assembly between *Ovtrpv1* and *Ovtrpv2* into an active heteromeric channel that is directly gated by aversive substances (Figure 8). This finding is in line with previously characterised invertebrate TRPV receptors. In both *C. elegans* and *D. melanogaster*, co-expression of the pair *osm-9/ocr-2* (but more robustly of *osm-9/ocr-4*) and *Nan/Iav* respectively, produced dose-dependent responses to NAM (Upadhyay *et al*., 2016; Griffin *et al*., 2025). Behavioural and cell-studies revealed this substance is a noxious bitter cue in invertebrates, eliciting avoidance responses and leading to a cell-death inducing accumulation of NAM through desensitisation of TRPV channels (Upadhyay *et al*., 2016; Ishikawa *et al*., 2023). Our data are therefore consistent with a conserved TRPV role in invertebrate sensory detection and cell metabolic regulation that are worth pursuing in the future.

## Conclusions

Altogether, this work identified two functional TRPV channels in *O. vulgaris* and contributed to providing precise annotation and experimental validation of two important molecular determinants of nociception in cephalopods. Heterologous functional characterisation of these octopus TRPV channels in *C. elegans* and in the *Xenopus* oocyte recombinant system, suggests they are polymodal and may have a physiological role in the octopus response to aversive chemical and mechanical cues.

The sensory role of *Ovtrpvs* is supported by our endpoint PCR analysis of a wide set of tissues, demonstrating expression in sensory and nervous tissues. However, future analysis should focus on qPCR and *in situ* hybridisation to confirm the potential selective expression in the tip of the arm and sucker relative to other non-sensory tissues.

Finally, the established phylogenetic relationship, locating OvTRPV1 and OvTRPV2 in distinct evolutionary branches of the vanilloid TRP channel subfamily, as well as the recombinant experiments showing the requirement for co-assembly of OvTRPV1 and OvTRPV2 to generate a functional channel, provides a route to experimentally investigate the pathways and the modalities in which *Ovtrpvs* operate. At the same time it fosters the development of experimental platforms that might identify important environmental sensory cues. Recent advances have in fact shown octopus evolved species-specific receptors but that they also exploit conserved ones (van Giesen et al., 2020), showing the importance for cephalopods to recognise and classify stimuli in the environment through an elaborated sensory system able to detect, integrate and select against nociceptive cues.

## Materials and Methods

### In silico analysis

#### BLAST search criteria to identify *O. vulgaris* transcripts

The protein sequence of *Homo sapiens* TRPV1 (*Hstrpv1*) that encodes the human capsaicin receptor was retrieved from UniProtKB/Swiss-Prot (release 2021_04) and blasted against the previously published *O. vulgaris de novo* assembly transcriptome (Petrosino, 2015; Petrosino *et al*., 2022) using the TBLASTN algorithm (BLAST+ v2.10.0+) with a standard threshold of 10e^-5^ used as a cut-off. The reference sequences were also used in a BLASTp search against the predicted proteome obtained from the most recent sequenced *O. vulgaris* genome (Destanović *et al*., 2023), using the same parameters described above. The hits retrieved from both searches, were blasted and compared against *Octopus bimaculoides* (<u>ASM119413v2</u>), *Octopus sinensis* (<u>GCA_006345805.1</u>) and *Aplysia californica* (<u>PRJNA13635</u>) assemblies to facilitate assessment of completeness.

In the case of the *O. vulgaris* chemotactile receptor (CRTs) orthologues, *O. bimaculoides* published sequences were retrieved from NCBI and used for a BLASTp search against *O. vulgaris* genome using the parameters mentioned above (Supplementary Table S1). All the resulting candidates were then aligned against the *H. sapiens* acetylcholine receptor α7 subunit (NP_000737.1) prior selection, to check for the lack of the classical vicinal cysteines involved in the neurotransmitter binding, which signatures cephalopod CRTs (van Giesen *et al*., 2020).

#### Exon-intron boundaries detection

The reconstructed *Ovtrpv* transcript sequences were blasted against the assembled *O. vulgaris* genome (Destanović *et al*., 2023) to localise their chromosomal location. The closest match was downloaded and converted into a BLAST custom database through the ncbi/suite tool container in Docker (Merkel, 2014). Alignment between the *Ovtrpv1* and *Ovtrpv2* cDNA from *O. vulgaris* or other species (i.e. *O. sinensis*) and the genomic sequence was performed using ncbi/magicblast tool container (Boratyn *et al*., 2019). The results were sorted using biocontainers/samtools (Danecek *et al*., 2021) and the data were processed and visualised using Tablet software (The James Hutton Institute, v1.21.02.08, Milne *et al*., 2013).

#### Secondary and 3D structure prediction and modelling

The assembled contig cDNA was translated to its predicted amino acid sequence through Expasy translate (Gasteiger *et al*., 2003). Each octopus predicted protein of interest was analysed using DeepTMHMM −1.0 to obtain a prediction of the secondary structure (Hallgren *et al*., 2022). A 3D reconstruction of the proteins’ single subunit was obtained using AlphaFold 3 (Abramson *et al*., 2024) and their representation within the cell membrane was modelled through PPM3 Server (Lomize, Todd and Pogozheva, 2022). Analysis of the conserved protein functional domains was performed using NCBI conserved domain (Lu *et al*., 2020).

Alignment with other species orthologue proteins was carried out using Clustal Omega (Madeira *et al*., 2022) and identity and similarities among the sequences were analysed with EMBOSS Water Pairwise Sequence Alignment (Madeira *et al*., 2024) using a BLOSUM62 matrix.

#### Cluster analysis and phylogenetic investigation of OvTRPVs

Reference protein sequences for TRP channels from *H. sapiens, C. elegans and Drosophila melanogaster* as well as other cationic ion channel sequences such as voltage-gated sodium, potassium and calcium ion channels were used to perform a BLAST search (e-value 1e^-10^ and 40 maximum target sequences) against the complete proteome of 13 representative species from Mammalia, Cephalopoda, Gastropoda, Bivalvia, Insecta, Crustacea and Nematoda. This proteome selection was based on the best Benchmarking Universal Single-Copy Orthologs (BUSCO) value (Supplementary Table S2). All the results from each species were sorted and processed through CD-hit using a 0.95 similarity threshold to remove shorter isoforms or redundant sequences (Li and Godzik, 2006; Fu *et al*., 2012). The resulting unique sequences from all the species (1102 sequences) were then used as a query to perform a CLuster ANalysis of Sequences (CLANS) (Zimmermann *et al*., 2018) with the BLAST HSPs e-value of 1e^-12^ and BLOSUM62 scoring matrix (Supplementary File S1). Visualisation of the sequences relationship and cluster organisation was performed with the CLANS toolkit with a set p-value threshold of 1e^-60^ (Zimmermann *et al*., 2018). All the sequences of the cluster which included the target *Ovtrpv1* and *Ovtrpv2* sequences (50 sequences, Supplementary File S2) were selected and used to generate a phylogenetic tree. Alignment of the sequences was performed using MAFFT v7.526, E-INS-i method (Katoh and Standley, 2013). The results were manually curated to exclude poorly aligned sequences (2) and then trimmed with TrimAl (Supplementary File S3) using gappy-out mode and default parameters (Capella-Gutiérrez, Silla-Martínez and Gabaldón, 2009). The best-fit model of evolution was selected with ModelFinder (LG+I+G4 chosen according to the Bayesian Information Criterion) in IQ-TREE3 v3.0.1 (Wong *et al*., 2025). Tree branches were obtained using aLRT-SH with 1,000 replicates and ultrafast bootstrap method. The final phylogeny was visualised using FigTree v1.4.4 (http://tree.bio.ed.ac.uk/software/figtree/). The tree was rooted using the vertebrate TRPV channel subfamilies.

### Animal samples

#### *O. vulgaris* samples collection

Young adults of *O. vulgaris* were caught by local artisanal fishermen in the Bay of Naples, Italy and humanely killed for tissue collection. The following tissues were dissected and stored in 50 μL RNA later for subsequent analysis: supra-oesophageal mass (SEM), sub-oesophageal mass (SUB), optic lobe (OL), stellate ganglion (StG), gastric ganglion (GG), arm (muscle + axial nerve cord, at 50% of its length), tip of the arm (TIP), sucker (Su), intestine (Int), stomach (Stom), white body (WB), anterior salivary gland (ASG), posterior salivary gland (PSG), digestive gland (DG), haemocytes (Hc), kidney (Kid), branchial heart (BrH), skin, mantle (Man) and gill (Supplementary Table S3).

#### *C. elegans* strains and husbandry

The following nematode strains were utilised in this study: CX10 *osm-9 (ky10);* CX4544 *ocr-2 (ak47)*; the Bristol N2 was used as the wild-type (WT) strain. The strain genotype of the *osm-9 (ky10)* and *ocr-2 (ak47)* mutants was confirmed by designing a pair of primers flanking the region of mutations (Supplementary Table S4) and sequencing (Eurofins Genomics).

*C. elegans* were cultured and maintained as described in Brenner (1974). Three days prior to any assay, gravid adult worms were put on culture plates to lay eggs for 4h and then removed to produce a synchronised population to be tested at the L4+1-day old (young adult) stage. In the case of transgenic lines, L4 worms expressing GFP derived from co-marker plasmid were selected from the synchronised population and incubated overnight at 20 °C 24h prior the experiment.

### Molecular biology

#### RNA extraction and cDNA synthesis from *O. vulgaris* tissues

Small resections (7-40 mg) of the indicated frozen tissues were collected in 2 mL Eppendorf tubes containing 500 µl TRIzol® (Invitrogen™), snap frozen and homogenized using a Handheld Homogeniser SHM1 (Cole-Palmer). After 5 min at room temperature, 200 µl of chloroform (Sigma-Aldrich) were added, mixed, and incubated on ice for 15 min. Samples were centrifuged (12,000 x g, 4° C) and the total RNA upper aqueous layer purified using PureLink® RNA Mini Kit (Invitrogen^TM^). DNase I treatment (Invitrogen^TM^) was performed to remove potential contaminating genomic DNA. Quality (260/280 ≈ 2.0) and quantity of extracted RNA were assessed through UV-visible absorption measurements (NanoDrop™ 2000/2000c Spectrophotometers). For each sample, 1 µg of RNA was used for reverse transcription following the manufacturer’s protocol (SuperScript™ III Reverse Transcriptase, Invitrogen^TM^). The cDNA samples were stored at −20°C until further use. As previously reported, the mollusc ‘hidden break’ does not allow a sufficient gel resolution to assess the RNA quality due to the co-migration of the 18S and 28S rRNA fragments (Natsidis *et al*., 2019; Adema, 2021). Therefore, only the tissues from which we successfully amplified a control gene, *cullin 1 (cul1)*, were selected for further use in PCR amplification of the genes of interest (Supplementary Table S3).

#### Primer design and end point PCR

Primer sequences of the genes of interests were designed using ApE-A plasmid Editor© v3.1.3 software and purchased from Eurofins Genomics (Supplementary Table S4). Once diluted to a working concentration of 25 pmol/µL, the primers were used to amplify the sequence in the tissues described above using 1 µL of cDNA as template in a PCR Phusion™ High-Fidelity DNA Polymerase (Thermo Scientific™) following manufacturer’s instructions.

#### Promoter and genes synthesis for cloning

The promoter region of the *osm-9* gene (*Posm-9*, approximately 1.6 kb – Colbert, Smith and Bargmann, 1997), the *C. elegans osm-9* (*Celeosm-9*) and the *Hstrpv1* cDNA sequences were synthesised using the Integrated DNA Technologies Gene synthesis service (Integrated DNA Technologies, USA). The promoter region of the *ocr-2* gene (*Pocr-2*, approximately 2.5 kB-Sokolchik *et al*., 2005) and the *O. vulgaris trpv2* (*Ovtrpv2*) were synthesised using the Genscript gene synthesis service and subcloned into the pcDNA3.1 vector (Genscript Biotech, UK). The pcDNA3.0 plasmid containing the *C. elegans ocr-2* cDNA sequence (*Celeocr-2*) was a kind gift from Prof. Cornelia Bargmann (Rockefeller University, NY, USA). All the plasmid sequences received, synthesised and cloned were verified for their authenticity using the Oxford Nanopore Whole Plasmid Sequencing (WPS) service from Eurofins Genomics. A missense mutation (AAC to AGC) in *ocr-2* cDNA leading to N695S in the protein sequence was identified in the original *pcDNA3-ocr-2* plasmid. However, the cDNA still showed a successful behavioural rescue when expressed in *ocr-2 (ak47)* mutant worms suggesting it is functionally silent, consistently with previous investigations (Tobin *et al*., 2002; Ohnishi *et al*., 2020).

#### Cloning for expression in *C. elegans*

The *Pocr-2/Posm-9* sequences were cloned (using *HindIII/XhoI*) into the pWormgate2 vector (Johnson, Behm and Trowell, 2005) to produce a pDEST-*Pocr-2* or pDEST-*Posm-9* vector for suitable expression in *C. elegans*. The full-length transcript sequences of *Ovtrpv1*, Ov*trpv2* and *Celeocr-2* were amplified (Supplementary Table S4) using Phusion™ High-Fidelity DNA Polymerase (Thermo Scientific™) and the A’ overhangs were added using GoTaq® G2 DNA Polymerase (Promega; 30 min at 72 °C) prior to ligation into pCR™8/GW/TOPO™ TA (Invitrogen™). This sequence was then transferred into the pWormgate2, downstream of the corresponding promoter *via* attR/attL recombination using a Gateway™ LR Clonase™ II Enzyme mix (Invitrogen™). This produced the final constructs: pDEST-*Posm-9*::*Ovtrpv2*, pDEST-*Pocr-2*::*Ovtrpv1*, pDEST-*Pocr-2::Celeocr-2*. The *Celeosm-9* cDNA sequence was cloned via *KpnI/SacI* downstream of *Posm-9* to produce pDEST-*Posm-9*::*Celeosm-9* final construct (Supplementary File S4). The plasmids were transformed into chemically competent bacterial cells (Thermo Scientific™) and the amplified plasmids purified from overnight cultures. These were then purified using the Monarch® Spin Plasmid Miniprep Kit (New England Biolabs®, UK) and authenticated by sequencing (Eurofins Genomics).

#### Cloning for expression in *Xenopus laevis* oocytes

The sequences of *Ovtrpv1, Ovtrpv2,* were cloned into the multiple cloning site of the oocyte expression vector pTB207 *via HindIII/NotI*. The sequence of *Hstrpv1* was cloned via PCR amplification to introduce the compatible cutting sites *EcoRI-NotI* (Supplementary Table S4). The pTB207 vector was a kind gift from Jean-Louis Bessereau’s Lab (Claude Bernard University, Lyon, France). These plasmids were transformed into chemically competent bacterial cells (Thermo Scientific™) and the amplified plasmids purified from overnight cultures. These were then purified using the QIAGEN Plasmid Maxi Kit (QIAGEN) and authenticated by sequencing (Eurofins Genomics). The final construct sequences are available in Supplementary File S5.

#### C. *elegans* fosmids purification and validation

The fosmids B0212.3 *osm-9* (clone ID: WRM066bG12) and C07G1 *ocr-2* (cloneID: WRM0634bB10) were purchased from K.K. DNAFORM (Yokohama, Japan). The amplified fosmids were purified using the QIAGEN Plasmid Maxi Kit (QIAGEN). PCR verification was performed using primers designed to amplify an internal region of *osm-9*/*ocr-2* which was authenticated by sequencing (Eurofins Genomics). The construct sequences are available in Supplementary File S6.

#### Microinjections

The purified pWormgate2 plasmid (Johnson, Behm and Trowell, 2005) containing the indicated genes was microinjected (40 ng/µl of plasmid,10 ng/µl in the case of fosmid) and 30 ng/µl of *Pmyo-2::gfp* or *Pmyo-3::gfp* (Addgene). These co-injection markers highlight the pharyngeal or the body wall muscle respectively. The microinjection was performed following the protocol described in Mello et al. (1991) and Mello and Fire (1991) with aluminosilicate glass capillaries (1.0 mm OD, 0.78 mm ID, Harvard Apparatus). Injections were conducted as previously described and injected worms were transferred onto a new seeded plate (Calahorro *et al*., 2022). Progeny from the injected worm was selected based on the fluorescence and transferred onto separate seeded plates to establish stable transgenic lines for propagation (F2 generation). The following transgenic lines were obtained: *ocr-2 (ak47) Pmyo-3::gfp; ocr-2 (*WRM0634bB10) Pmyo-3::gfp; *pDEST-Pocr-2::Ovtrpv1 Pmyo-3::gfp; pDEST-Pocr-2::Celeocr-2 Pmyo-3::gfp* in *ocr-2(ak47)* mutant background; *osm-9(ky10) Pmyo-3::gfp; osm-9 (WRM066bG12) Pmyo-2::gfp; pDEST-Posm-9::Celeosm-9,Pmyo-3::gfp, pDEST-Posm-9::Ovtrpv2* in *osm-9 (ky10)* mutant background.

### Behavioural assays

#### Drop assay to test chemoaversion in *C. elegans*

The behavioural analyses were conducted in experimental arenas in which a microscope digital camera (MU1403B, AmScope^TM^) was mounted and worm behaviour in response to the indicated treatments was recorded. The captured videos were analysed through manual scoring using AmScope Software v.10.11.2024. The assays were time stamped by the digital camera and the speed and quality of the responses assessed using the criteria listed below. The experimenter and observer were blind to the genotype of the strains under investigation.

Chemical aversion in *C. elegans* was performed using an adapted version of the classical acute drop assay (Hilliard, Bargmann and Bazzicalupo, 2002; Hilliard *et al*., 2004). Ten L4+1-day old worms from indicated lines were transferred onto a 9 cm unseeded NGM plate and left undisturbed for at least 20 min. This allows the worms to transition to roaming behaviour encompassing periods of extended forward runs (Gray, Hill and Bargmann, 2005). A small drop of noxious cue was delivered in front of a moving worm through a small glass capillary (1.0 mm OD, 0.78 mm ID, Harvard Apparatus) attached to a syringe. A binary score was assigned with a positive response (1) or negative response (0) for each compound tested within 5 s of exposure to the cue and was based on the average number of reversals being equal or higher to the average number showed by N2s. The worms for each condition were tested by exposing them to the drop only once. To trigger low pH response, acetic acid (CH_3_COOH; Fisher chemical) was dissolved in M9 buffer at a final pH of 3 (M9, pH 3). M9 buffer (pH 7) was checked not to cause any response to N2s when administered alone.

#### Nose touch assay

Briefly, an eyebrow was placed perpendicular to the front of a moving L4+1-day old worm until the animal hit the hair (Kaplan and Horvitz, 1993). Worms were given a binary score indicating nose touch sensitivity (1) or insensitivity (0). The sensitive worms halted or/and reversed following the collision. The insensitive worms kept moving forward and tried to climb or cross the hair. The experiment was performed in a single set of 10 consecutive touches on 5 worms per strain. The final ratio of sensitive/insensitive responses per worm was calculated and compared between N2, mutant and transgenic strains.

#### Volatile aversive response

Following the same husbandry described for the drop assay, 10 worms expressing long forward runs were individually exposed to 30% 1-octanol (Sigma-Aldrich) solution only once. A thin platinum wire (Ø 0.1 mm, Agar Scientific) was dipped into the solution and then waived in front of a moving worm until the animal stopped and initiated a backward movement. The average latency to start a reversal (s) was recorded with a cut-off of 15 s.

#### Data analysis and statistics

In the behavioural assays the transgenic lines that showed a performance that reached two standard deviations from the mean of the mutant line were considered rescued and included in the analysis.

Data were analysed using either one-way or two-way parametric analysis of variance (ANOVA). Post-hoc comparisons were performed using the Dunnett’s multiple comparisons test. A level of probability set at p<0.05 was used as statistically significant. Statistics were performed with GraphPad Prism version 10 for Windows (GraphPad Software, Boston, Massachusetts, USA).

### TRPVs ligand activation

#### Electrophysiology of *Xenopus* Oocytes

Defolliculated *Xenopus laevis* oocytes were obtained from EcoCyte Bioscience and maintained in ND96 solution as follows (in mM): 96 NaCl, 1 MgCl _2_, 5 HEPES, 1.8 CaCl _2_, and 2 KCl adjusted to pH 7.4. The plasmids of interests, *pTB207-Hstrpv1, pTB207-Ovtrpv1, pTB207-Ovtrpv2,* were linearised using *PacI*, followed by DNA purification (Zymo) and elution in RNAase free water. cRNA was synthesised using the T7 mMessage mMachine transcription kit (Thermo Fisher Scientific) according to the manufacturer’s protocol (with incubation for 6h at 37 °C). RNA was purified using the GeneJET RNA purification kit (Thermo Fisher Scientific) and quantified using a NanoDrop spectrophotometer. Oocytes were injected with 50 nl of 500 ng/µl RNA individually or co-expressed (*pTB207-Ovtrpv1/pTB207-Ovtrpv2*) using the Roboinject system (Multi Channel Systems). Injected oocytes were incubated at 18 °C in ND96 solution until the day of recording, 4 days post-injection.

Two-electrode voltage-clamp (TEVC) recordings were conducted using the Roboocyte2 System (Multi Channel Systems). Measuring head electrode resistance was approximately 400-1200 kΩ, pulled on a P-97 Micropipette Puller (Sutter Instrument). Electrodes contained AgCl wires backfilled with a 1 M KCl and 1.5 M KAc mixture. Oocytes were clamped at −60 mV during continuous recording at 500 Hz. Compounds applications (capsaicin, nicotinamide, ND96) lasted for 20 s for capsaicin followed by a 60 s wash with ND96, and 600 s for nicotinamide followed by a 200 s or 1200 s wash with ND96. Perfusion speed was set to approximately 3 ml/min throughout. Uninjected oocytes were also tested and did not respond to any of the compound tested. At least 7 oocytes were tested for each condition. Data were analysed using the Roboocyte2+ software.

#### Compounds and stimuli tested

The following compounds were utilised in this study: capsaicin natural (Biosynth), acetic acid glacial (FisherChemical), 1-octanol (Sigma-Aldrich), nicotinamide (NAM, Sigma-Aldrich). Stock solutions were prepared either in ethanol or DMSO/distilled water, and the final working concentration was reached by dissolving the solution in ND96 medium prior to performing the assay. Saline-injected oocytes did not respond in any case tested.

### Declarations

#### Ethics approval

*O. vulgaris* specimens were manipulated for the sole scope of tissue collection following the local Animal Welfare Body authorisation (Ethical Clearance: ecACR-2302ts36). Animals were humanely killed adopting the principles described in Annex IV of Directive 2010/63/EU and following recommendations from Andrews *et al*. (2013), Fiorito *et al*. (2015), Butler-Struben *et al*. (2018) to ensure responsible and ethical use of animal-derived materials and to adhere to the principles of Replacement, Reduction, and Refinement (3Rs). The tissues were collected by a FELASA certified (function D) competent person. All the experiments have been carried out in compliance with the Ethics and Research Governance Online II (ERGO II) policy (nr 79739) in place at the University of Southampton.

## Consent for publication

Not applicable

## Availability of data and materials

The datasets (Supplementary Tables S1-S4 and Supplementary Files S1-S6) supporting the conclusions of this article are available in the Zenodo repository at the following link: 10.5281/zenodo.18377710.

## Authors’ contributions

EMP: conceptualisation, data curation, *in silico* analysis, experimental investigation, methodology, validation, visualization and writing–original draft

HB: conceptualisation, data curation, experimental investigation, methodology, visualization and writing– review and editing, resources

VOC: conceptualisation, methodology, writing–original draft, writing–review and editing, funding acquisition and supervision, resources

LHD: conceptualisation, methodology, writing–original draft, writing–review and editing, funding acquisition and supervision, resources

LAYG: methodology, validation, visualization and writing– review and editing, resources PI: writing–review and editing, resources

GF: conceptualisation, writing–review and editing, funding acquisition

JD: conceptualisation, methodology, writing–original draft, writing–review and editing, funding acquisition and supervision, resources

## Acknowledgements

Strains were provided by the Caenorhabditis Genetic Center (CGC), funded by NIH Office of Research Infrastructure Programs (P40 OD010440). We thank Prof Cori Bargmann for kindly providing us with the pcDNA3.1-*ocr-2* construct. We thank Jean-Louis Bessereau’s Lab (Claude Bernard University, Lyon, France) for kindly providing the pTB207 vector. We thank Dr Iris Hardege and Tom Reynoldson for advice and support with electrophysiology. We thank Eng Marco Pieroni who kindly helped with the setup and use of the described Docker containers.

